# Matrix tropism influences endometriotic cell attachment patterns

**DOI:** 10.1101/2025.02.22.639314

**Authors:** Hannah S. Theriault, Hannah R.C. Kimmel, Alison C. Nunes, Allison Paxhia, Sarah Hashim, Kathryn B.H. Clancy, Gregory H. Underhill, Brendan A.C. Harley

**Affiliations:** Dept. of Bioengineering; Carl R. Woese Institute for Genomic Biology; Dept. Chemical and Biomolecular Engineering; Dept. of Anthropology; Dept. of Women & Gender Studies; Center for Gender and Sex in Health; Cancer Center at Illinois University of Illinois at Urbana-Champaign Urbana, IL 61801

**Keywords:** Endometriosis, lesion initiation, extracellular matrix, tissue engineering

## Abstract

Due to the extended period for clinical diagnosis, the etiology of endometriotic lesion initiation is not well understood or characterized. Endometriotic lesions are most often found on pelvic tissues and organs, especially the ovaries. To investigate the role of tissue tropism on ovarian endometrioma initiation, we adapted a well-characterized polyacrylamide microarray system to investigate the role of tissue-specific extracellular matrix and adhesion motifs on endometriotic cell attachment, morphology, and size. We report the influence of cell origin (endometriotic vs. non-endometriotic), substrate stiffness mimicking aging and fibrosis, and the role of multicellular (epithelial-stromal) cohorts on cell attachment patterns. We identify multiple ovarian-specific attachment motifs that significantly increase endometriotic (vs. non-endometriotic) cell cohort attachment that could be implicated in early disease etiology.

## 1. Introduction

Endometriosis is an estrogen-dependent, chronic-inflammatory disease impacting approximately 5-10% people who menstruate [1, 2] and up to 35-50% of menstruators experiencing infertility [3, 4]. Endometriosis is histologically defined by the presence of endometrial-like tissues on ectopic, non-native, locations [5] that signal with, attach to, and colonize pelvic tissues and organs. One of the most common lesion locations is on the ovaries, referred to as endometriomas. These lesions undergo cyclic proliferation and degradation over the course of the menstrual cycle akin to native endometrial tissues [5]. Their presence can result in incapacitating pelvic pain, dysmenorrhea, menorrhagia, among other symptoms [6, 7]. Diagnosis is often delayed, taking an average of 4-11 years after symptom onset [6]. This gap contributes to our poor understanding of lesion initiation. One widely accepted etiological theory involves retrograde menstruation, where menstrual effluent is refluxed into the pelvic cavity [8]. However, this occurs in nearly 90% of menstruators [9], suggesting retrograde menstruation is not sufficient alone for lesion formation. Other theories include metaplasia of peritoneal cells, defective embryogenesis, oxidative stress, or immunological factors [5, 10–15], but no singular theory explains the idiosyncratic distribution of lesions. We hypothesize extracellular matrix (ECM) tropism provides a lens to investigate processes underlying lesion initiation and invasion *in vitro*.

The ECM contains a range of collagens, laminins, fibronectin, proteoglycans and glycosaminoglycans (GAGs) [16]. Irregular ECM deposition and remodeling patterns have been implicated in cancer invasion, metastatic potential, and progression [17–19] as well as endometriosis progression [20]. The ovarian ECM microenvironment varies based on depth (with a stiffer external cortex and a softer inner medulla) [21, 22], a function of age, and disease status (1.79 ± 0.08 kPa to 4.56 ± 2.03 kPa) [23, 24]. Due to similarities in processes shared by endometriomas and ovarian cancer, we hypothesize that lesion initiation may be significantly influenced by cell-matrix (e.g., integrin) or cell-cell (e.g., cadherin) adhesion molecules in the ovarian microenvironment [25–29]. While cell adhesion strength is positively correlated with time allowed for attachment, early interactions may be critically dependent on specific integrin and cell adhesion activation [25, 30]. Studies of cancer progression often consider the role of the unique local tissue microenvironment (e.g., tissue tropism) [31–33], motivating our effort to consider reciprocal relationships between endometriotic cells and the local tissue microenvironment. Improved understanding of the collective influence of mechanically driven cell-cell and cell-matrix interactions within the ovarian microenvironment on endometriotic cell adhesion will provide a new model to understand early stages of endometrioma lesion initiation.

Endometriotic lesions are multicellular and contain stromal cells with smaller fractions of epithelial (or epithelial-like) cells, lymphocytes, myeloid-lineage cells and endothelial cells [34]. Previous models have begun to consider the activity of cellular components of endometriotic lesions such as the comparison of endometriotic epithelial-like cell lines in 3D spheroid culture examining cellular invasion capacity, epithelial to mesenchymal transition (EMT) phenotype, and cytokeratin and vimentin expression [35]. More recent models considered interactions between endometriotic lesion derived epithelial and stromal cell cohorts, finding the potential for cell cohorts to self-assemble into multilayered spheroids with an enhanced invasive capacity [36]. While these models are increasingly replicating cellular aspects of endometriotic lesions, they are commonly performed in simplified biomaterial (e.g., Matrigel) that exhibit batch-to-batch variability and poorly defined matrix and biomolecular content. To highlight the matrix environment, we aim to identify key elements of the ovarian ECM that drive endometriotic cell attachment.

In this study, we adapted a well-characterized, high-throughput 2D microarray system [37–41] to investigate the influence of ECM composition, cell adhesion molecules, and stiffness on the adhesion and morphology of epithelial cells, stromal cells, and multicellular. We used a recently immortalized endometriotic stromal cell line (iEC-ESCs) [42] as well as an endometriotic epithelial-like cell line (12z) [43] to represent the primary components of endometriomas. We then compared non-endometriotic endometrial-derived endometrial epithelial cells (EECs) and endometrial stromal cells (ESCs). We describe these cells as endometriotic epithelial (EndoE), endometriotic stromal (EndoS), non-endometriotic epithelial (NonEndoE), and non-endometriotic stromal (NonEndoS). We identify shifts in cell adhesion and morphology in response to ovarian-specific matrix and cell adhesion factors, comparing results to control (liver) tissue factors where lesions are rarely observed [44]. This work demonstrates an integrated approach to consider the role of both matrix and cell adhesion tropism signals. Further, we explicitly consider the collective adhesion behavior of cohorts of EndoE and EndoS cells necessary for lesion initiation. Collectively, we report new technical capacities to consider multicellular adhesion as well as critical new insight that tissue tropism and the collective activity of endometriotic cell cohorts play an important role in shaping endometrioma lesion initiation and persistence.

## 2. Results

### 2.1. Microarrays support the addition of cell adhesion molecules and cadherins

A detailed graphical abstract of our experimental design is provided in **Figure 1**. We fabricated polyacrylamide (PA) microarrays with defined mechanical stiffness that contained pairwise combinations of cell adhesion molecules (CAMs), cadherins, and ECM biomolecules identified as ovarian-specific, liver-specific, or common (non-specific) (**Table 1 & 2**). When CAMs and cadherins were incorporated (**Fig. 2a**), there was limited cell adhesion to CAM-only or cadherin-only islands, however the addition of a CAM or cadherin to another ECM biomolecules significantly altered the cellular adhesion compared to the ECM biomolecule alone (**Fig. 2b & 2c**). As a result, we screened endometriotic cell attachment using microarrays containing 86 pairwise combinations of ECM biomolecules and CAMs or cadherins (**Table 1 & 2**). We quantified the number of attached cells of each type to each of the 86 combinations on substrates that presented physiological (1 kPa) or pathological/aged (6 kPa) stiffness (**Fig. 2d**). We identified a down-selected subset of islands (42 combinations) that supported an average adhesion of n ≥ 25 cells per island for endometriotic epithelial and stromal cell populations (**Fig. S1c**). Notably, this down-selected list of cell/matrix adhesion combinations had a greater frequency of ovarian (ovary-specific and common) combinations (79.1% of full list to 85.7% in the down-selected list) and a smaller frequency of liver-specific islands than the full list (20.9% to 14.3%) (**Fig. 2e & Table 2**).

**Figure 1:**
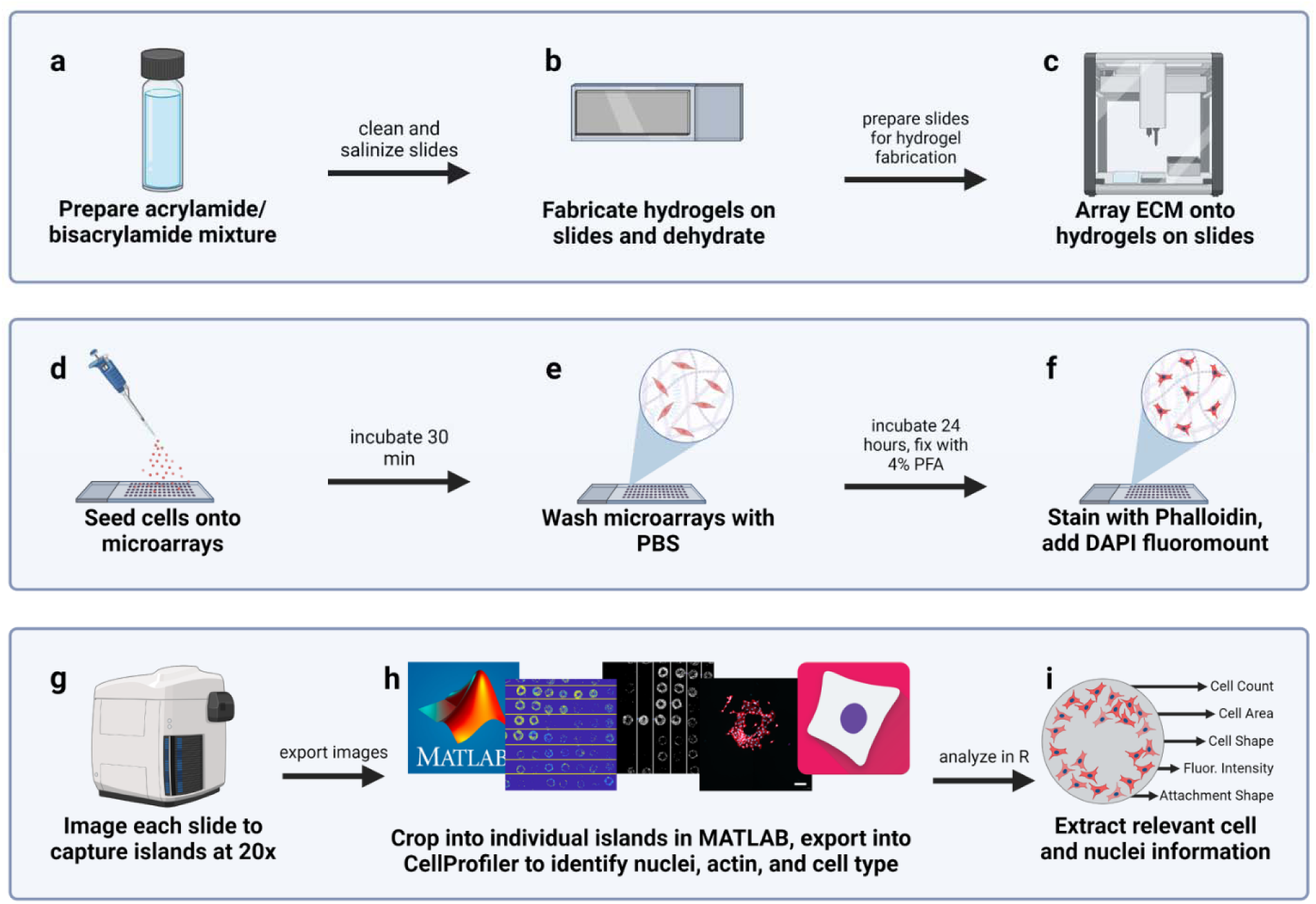
Methodology for preparing, seeding, and analyzing microarrays. a) 1 kPa and 6kPa PA solutions are prepared as described in 2.2. b) Slides are washed, etched, and silanized before prepolymer solution is deposited in the printing region of interest and cured and dehydrated. c) ECM combinations are diluted in printing buffer to a final concentration of 250 μg/mL and arrayed onto the slide(s) with a liquid handler. d) 200,000 cells per microarray were seeded in monoculture or coculture and incubated for 30 minutes. e) Microarrays were washed with PBS and then incubated for 24 hours at which point f) microarrays were fixed in 4% PFA and stained with phalloidin and DAPI with fluoromount. g) Fixed and mounted slides were imaged on the Zeiss AxioScan.Z1. h) Images were exported and cropped into identified islands using MATLAB. CellProfiler was used to identify fluorescent regions of the cell (nuclei, actin) and partition information including cell count, cell area, etc. i) The outputs of cell profiler are then analyzed in R Studio.

**Table 1:** Description of single ECM and cell attachment factors used in large microarray printing.

| Tissue Type | Biomolecule | Vendor | Catalogue # | Concentration |
| --- | --- | --- | --- | --- |
| Ovary | Collagen IV [50, 73] | Corning | 354233 | 1 mg/mL |
|  | Hyaluronan [74] | Lifecore Biomedical | HA60K-1 | 1 mg/mL |
|  | Syndecan-1 [74] | R&D Systems | 2780-SD-050 | 0.5 mg/mL |
|  | Glypican-1 [74] | R&D Systems | 4519-GP-050 | 0.5 mg/mL |
|  | Versican [74] | R&D Systems | 3054-VN-050 | 0.5 mg/mL |
|  | VCAM-1 [75] | R&D Systems | 862-VC-100 | 1 mg/mL |
|  | ICAM-1 [76] | R&D Systems | ADP4-050 | 1 mg/mL |
|  | EP-CAM [77] | R&D Systems | 960-EP-050 | 1 mg/mL |
| Liver | Collagen V [78] | Rockland | 009-001-107 | 1 mg/mL |
|  | Elastin | Millipore Sigma | E6902-1MG | 1 mg/mL |
|  | Tenascin C | R&D Systems | 3358-TC-050 | 0.5 mg/mL |
| Common | Collagen I [73, 74, 79] | EMD Millipore | 08-115 | 1 mg/mL |
|  | Collagen III [74, 79] | EMD Millipore | CC054 | 1 mg/mL |
|  | Fibronectin [73, 74, 79] | EMD Millipore | FC010 | 1 mg/mL |
|  | Laminin [73, 74, 79, 80] | Invitrogen | 23017-015 | 1 mg/mL |
|  | E-Cadherin [81] | R&D Systems | 8505-EC-050 | 1 mg/mL |

**Table 2:** Named and identified combinations of attachment biomolecules used in down-selected printings.

| Group | Biomolecule 1 | Biomolecule 2 | Abbreviation |
| --- | --- | --- | --- |
| Ovary | Collagen III | Collagen IV | C3:C4 |
|  | Collagen IV | Collagen I | C4:C1 |
|  | Collagen IV | Collagen IV | C4 |
|  | Collagen IV | Fibronectin | C4:FN |
|  | Collagen IV | Laminin | C4:LN |
|  | E-cadherin | Collagen I | ECAD:C1 |
|  | E-cadherin | Collagen III | ECAD:C3 |
|  | E-cadherin | Collagen IV | ECAD:C4 |
|  | EP-CAM | Collagen I | EPCAM:C1 |
|  | EP-CAM | Collagen III | EPCAM:C3 |
|  | EP-CAM | Collagen IV | EPCAM:C4 |
|  | Glypican-1 | Collagen I | GLY1:C1 |
|  | Glypican-1 | Collagen III | GLY1:C3 |
|  | Glypican-1 | Collagen IV | GLY1:C4 |
|  | Hyaluronan | Collagen I | HA:C1 |
|  | Hyaluronan | Collagen III | HA:C3 |
|  | Hyaluronan | Collagen IV | HA:C4 |
|  | Hyaluronan | Hyaluronan | HA |
|  | ICAM-1 | Collagen I | ICAM:C1 |
|  | ICAM-1 | Collagen III | ICAM:C3 |
|  | ICAM-1 | Collagen IV | ICAM:C4 |
|  | Syndecan-1 | Collagen III | SYN1:C3 |
|  | Syndecan-1 | Collagen IV | SYN1:C4 |
|  | VCAM-1 | Collagen I | VCAM:C1 |
|  | VCAM-1 | Collagen III | VCAM:C3 |
|  | VCAM-1 | Collagen IV | VCAM:C4 |
|  | VCAM-1 | Laminin | VCAM:LN |
| Liver | Collagen V | Collagen I | C5:C1 |
|  | Collagen V | Collagen III | C5:C3 |
|  | Elastin | Collagen I | ELN:C1 |
|  | Elastin | Collagen III | ELN:C3 |
|  | Tenascin-C | Collagen I | TC:C1 |
|  | Tenascin-C | Collagen III | TC:C3 |
| Common | Collagen I | Collagen I | C1 |
|  | Collagen III | Collagen I | C3:C1 |
|  | Collagen III | Collagen III | C3 |
|  | Collagen III | Fibronectin | C3:FN |
|  | Collagen III | Laminin | C3:LN |
|  | Fibronectin | Collagen I | FN:C1 |
|  | Fibronectin | Fibronectin | FN |
|  | Laminin | Collagen I | LN:C1 |
|  | Laminin | Fibronectin | LN:FN |

**Figure 2.**
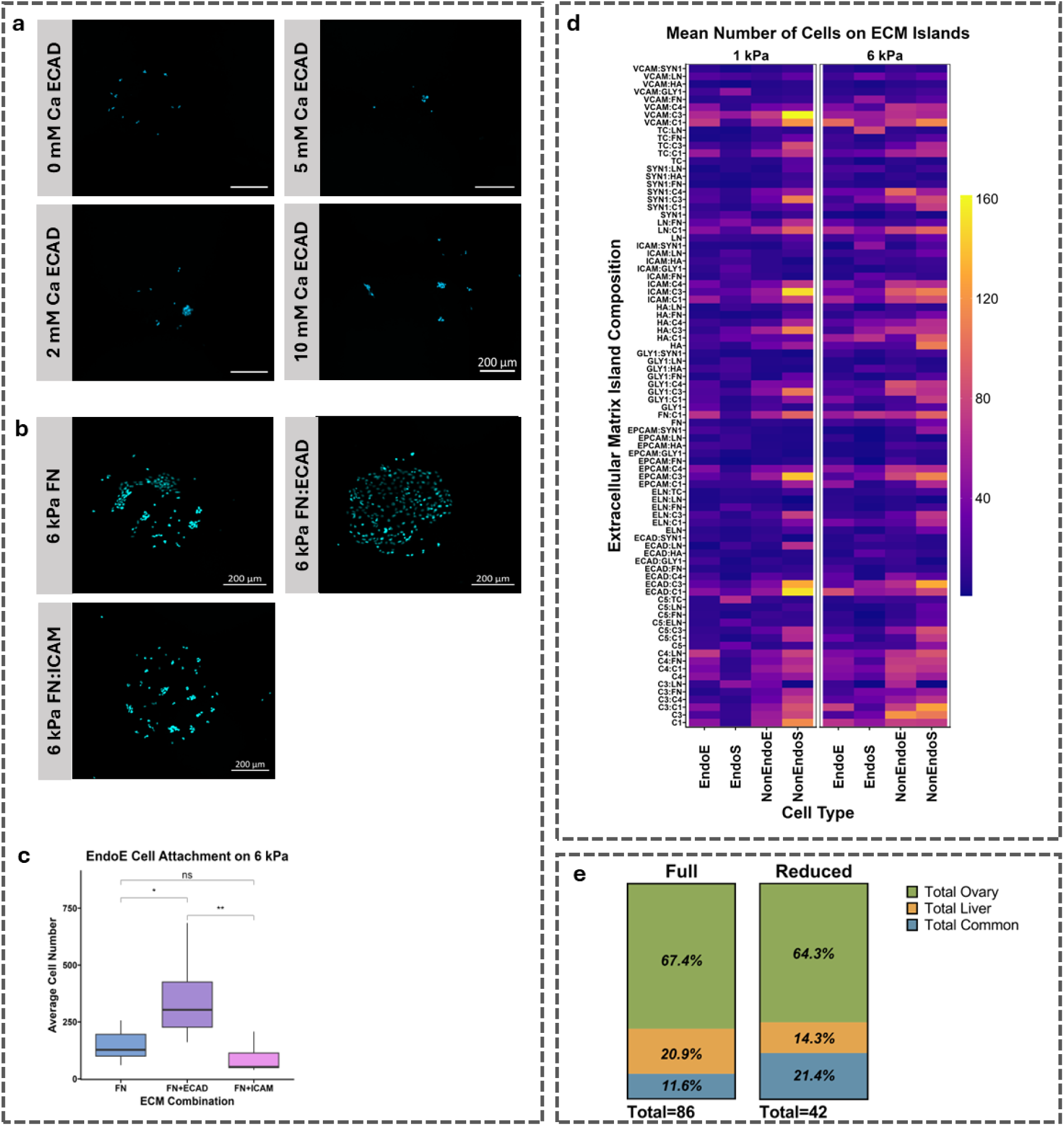
Examining the ability of PA microarrays to support the addition of CAMs and cadherins. a) Representative images of the early studies including CAMs and Cadherins showing increased cellular attachment with increased concentration of Calcium on Cadherin islands. 6kPa islands show low adhesion no calcium, 2.5 mM, 5mM, 10mM. b) Representative images of FN islands alone, with ECAD or with ICAM. c) Example callout of 12z endometriotic epithelial attachment comparing fibronectin alone, fibronectin with E-cadherin, and fibronectin with ICAM showing significantly increased attachment. d) The mean number of attached cells on the full array, each row represents an ECM Microenvironment, each column is a cell type faceted by stiffness. We can see that the adhesion profile varies within a column (cell type) and that there are some rows (ECMs) that have different adhesion levels for cell types or between stiffnesses. e) Visualization of the change in categorization contribution to the total classification of each island. Scale bar indicates 200 μm.

### 2.2. ECM composition plays a role in cell shape and area

We subsequently quantified the eccentricity (shape) and spread areas of endometriotic vs. non-endometriotic stromal and epithelial cells on the down-selected adhesive islands after 24 hours of culture via visualization of their actin structure (**Fig. 3a**). We hypothesized disease state and cell type would influence cell morphology and enable the ability to distinguish cell types in subsequent experiments. Eccentricity reports the ratio of elliptical foci and its major axis length that characterizes the spread cells, ranging from 0 (round) to 1 (elongated) (**Fig. 3b & S2a**). Statistically significant differences were observed in average eccentricities (across all adhesive island combinations and stiffnesses) between cell types (**Table S1).** Notably, stromal cells were significantly more elongated than the epithelial cells, with NonEndoE cells being the most rounded (**Fig. 3c**). We also observed significant differences in cell area as a function of adhesive combinations (**Table S1, Fig. 3d**). Stromal cells displayed a significantly larger area than epithelial cells, with EndoS cells having the greatest area and EndoE cells having the smallest. EndoE cells were significantly smaller than NonEndoE, while endometriotic stromal cells were significantly larger than non-endometriotic stromal cells (**Fig. 3e**). Key findings of area differences between stromal and epithelial cell types impacted the number of cells on islands which led us to develop a more robust comparison method utilizing z-scores (**Fig. S3a**).

**Figure 3.**
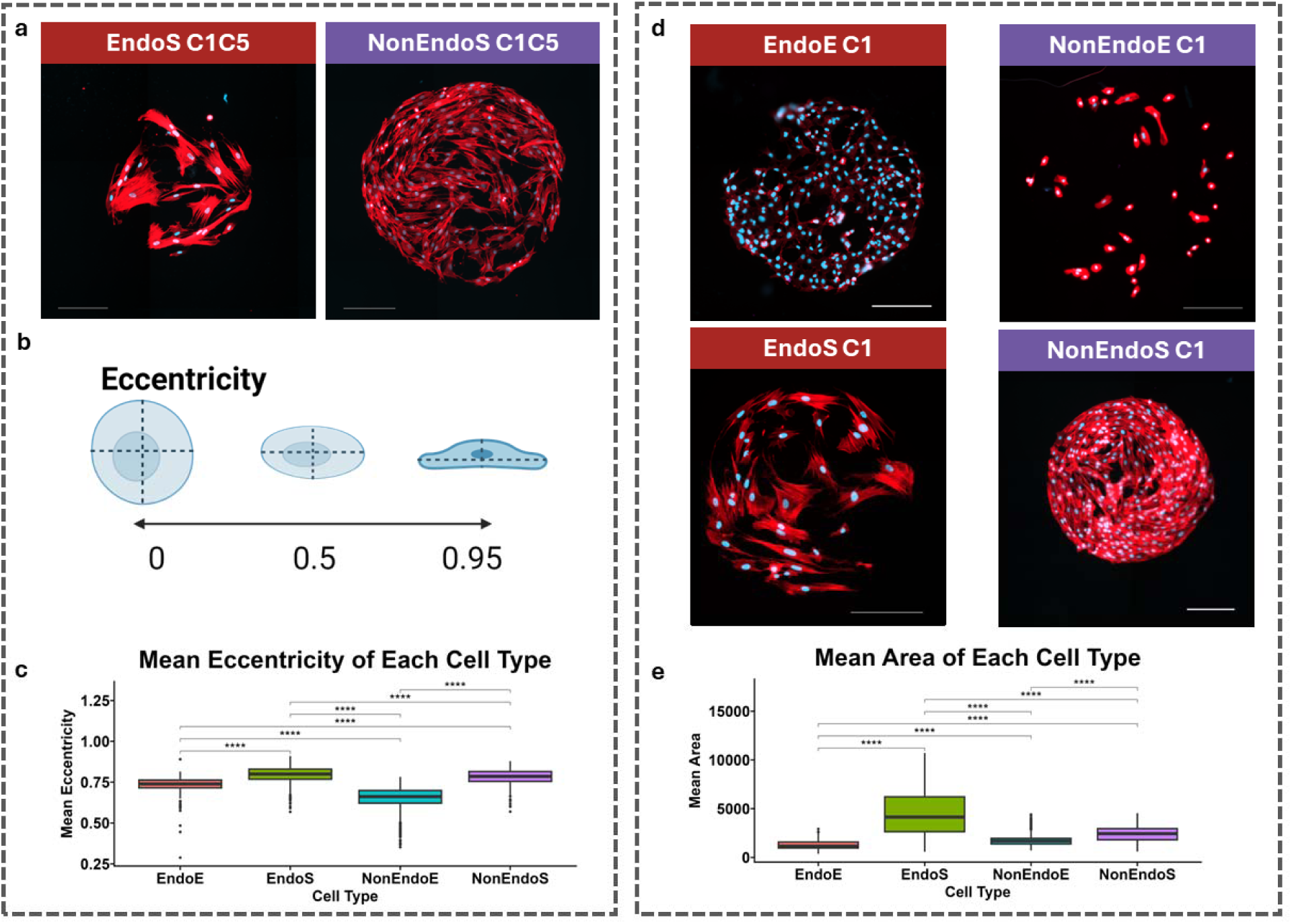
Observed changes in eccentricity by altering substrate stiffness and ECM. a) Representative images of visualization of stromal cell eccentricity differences. b) Eccentricity summary with zero being perfectly rounded and 1 being a straight line. Made in Biorender. c) Quantified general eccentricity of the four cell types. d) Representative images of all four cell types displaying major size differences. e) Quantified general cell area of the four cell types. Eccentricity and area are displayed here as an average over ECM conditions and stiffness. Scale bar indicates 200 μm.

### 2.3. Matrix tropism informs epithelial cell attachment

We then examined the effect of adhesive island properties on EndoE vs. NonEndoE cell attachment (**Fig. 4a**). Across all adhesion combinations, EndoE cell attachment was significantly increased on stiffer (6 kPa) vs. soft (1 kPa) islands while island stiffness did not significantly influence NonEndoE cells (**Table S2**, **Fig. 4b**). We then compared cell attachment, averaged over stiffness, between each adhesive island combination. We observed more significant variation in attachment for EndoE than in NonEndoE cells (Wilcox; gray: non-significant; purple: p ≤ 0.01; yellow: p ≤ 0.05) (**Fig. 4c**), suggesting EndoE cells are more sensitive to changes in adhesive island composition. We compared attachment between epithelial cell types (**Fig. S4a-c**), identifying 9 significant adhesive combinations at 1 kPa (7 combinations significantly increased EndoE cell attachment) and 20 significant adhesive combinations at 6 kPa (9 of which favored EndoE cell attachment). Adhesive combinations that increased EndoE cell (vs. NonEndoE) attachment at 1 kPa stiffness were primarily ovary-specific (C3:C4, C4, C4:C1, C4:FN, and VCAM:LN) and common (C3:C1 and FN). We also identified adhesive combinations at 6kPa that increased EndoE cell (vs. NonEndoE) attachment, the majority of which were either common (C1, C3:FN, FN, FN:C1, LN:C1 and LN:FN) or ovary-specific (C3:C4, C4:C1, and C4:FN) motifs (**Fig. 4d & Table S2**).

**Figure 4:**
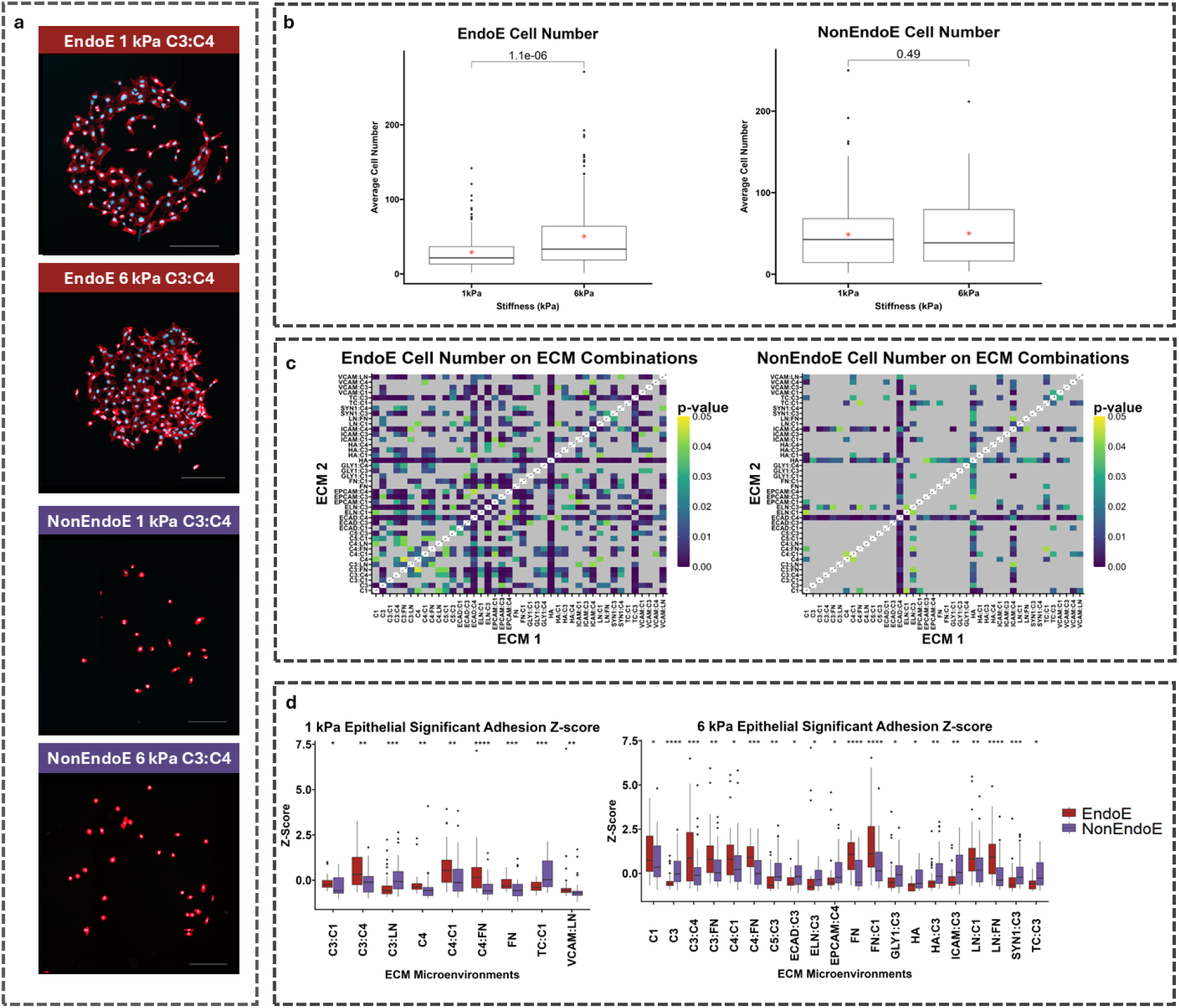
Reduced Arrays Comparison of Epithelial Cell Attachment. a) Representative images of endometriotic and non-endometriotic epithelial cells attached to C3:C4 islands across both stiffnesses. b) Quantification of the impact of stiffness on attachment of both epithelial cells displayed as the average across all ECM conditions for each cell type. c) Comparison of each ECM combination within each epithelial cell type to one another. Heatmaps display significant differences in color, self-comparisons in white, and non-significant comparisons in gray. d) Significant Z-score comparisons of EndoE and NonEndoE cells across both stiffnesses. Scale bar indicates 200 μm.

### 2.4. Matrix tropism informs stromal cell attachment

As lesions are multicellular, we next examined the adhesive biomolecule impact on endometriotic vs. non endometriotic stromal cell populations (**Fig. 5a**). Stiffness did not have a significant impact on cell attachment for both the endometriotic and non-endometriotic stromal cells (**Fig. 5b**). We subsequently compared attachment, averaged over stiffness between island combinations (**Fig. 5c**). As with the epithelial comparisons, there was found to be more variation between average adhesion to ECMs in the EndoS cells than the NonEndoS cells, indicating that EndoS cells are more sensitive to their microenvironment than their non-endometriotic derived complement. We identified attachment trends between EndoS and NonEndoS cells (**Fig. S5a-c**). Notably, significantly increased attachment of endometriotic to non-endometriotic stromal cells was observed in 6 of the 9 statistically significant comparisons on 1 kPa substrates, and 6 of the 10 statistically significant comparisons on 6 kPa substrates. The significant comparisons with increased endometriotic adhesion on the 1 kPa substrate were predominantly ovary-specific (C3:C4, C4:C1, ECAD:C4, HA, and SYN1:C4) with one liver-specific combination (TC:C3) and on the 6 kPa substrate were equally distributed between common (C3:FN, FN:C1 and LN:C1) and ovary-specific (ECAD:C4, HA, and SYN1:C4) conditions (**Fig. 5d, Table S3**).

**Figure 5:**
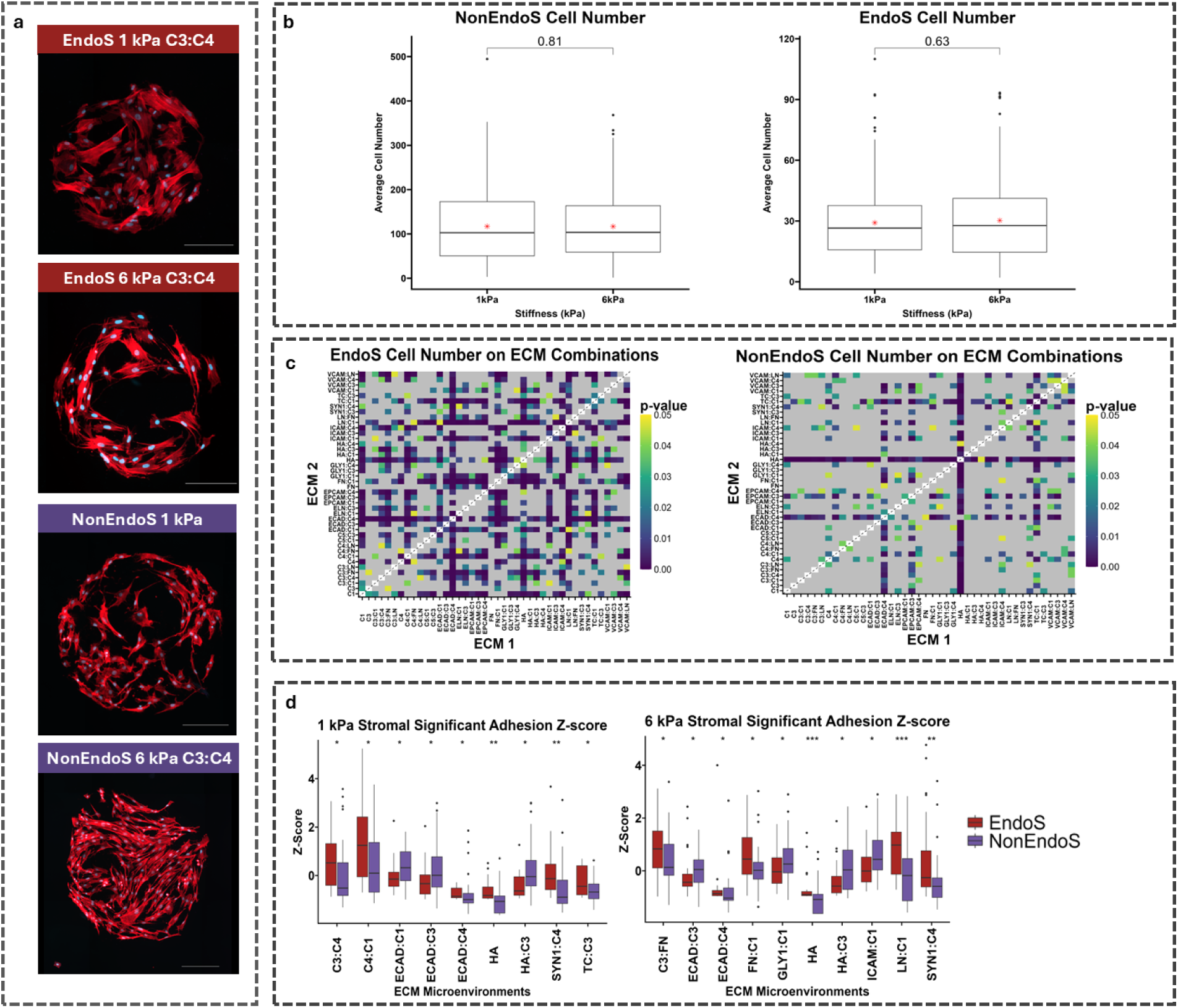
Reduced Arrays Comparison of Stromal Cell Attachment. a) Representative images of endometriotic and non-endometriotic stromal cells attached to C3:C4 islands across both stiffnesses. b) Quantification of the impact of stiffness on attachment of both stromal cells displayed as the average across all ECM conditions for each cell type. c) Comparison of each ECM combination within each stromal cell type to one another. Heatmaps display significant differences in color, self-comparisons in white, and non-significant comparisons in gray. d) Significant Z-score comparisons of EndoS and NonEndoS cells across both stiffnesses. Scale bar indicates 200 μm.

### 2.5. Coculture influences tissue tropism and cell morphology

We subsequently aimed to define the impact of the microenvironment on stromal-epithelial cell interactions in the context of lesion initiation. We seeded mixtures of stromal and epithelial cells onto the microarrays to create endometriotic and non-endometriotic cohorts. We used a physiologically relevant ratio of 75% stromal cells: 25% epithelial cells for both cohorts [34, 45]. To distinguish between the cell types for analysis, Cell-Tracker dyes were used to pre-label the stromal cells with green and epithelial cells with orange.

Notably, stiffer substrates increased attachment in co-cultures compared to softer substrates for all cell types (**Fig. S6i-j**). We observed significantly higher eccentricity of endometriotic epithelial cells in co-culture at 6 kPa compared to monoculture (**Fig. 6a**). Endometriotic stromal cells also exhibited differential eccentricity on the 1 kPa condition, becoming less eccentric in co-culture (**Fig. 6a**). We did not see distinct patterns with endometriotic epithelial cell area between co-culture and monoculture, however, we did observe endometriotic stromal cells on both stiffness conditions were smaller in co-culture than in monoculture (**Fig. 6b & S6d-h**).

**Figure 6:**
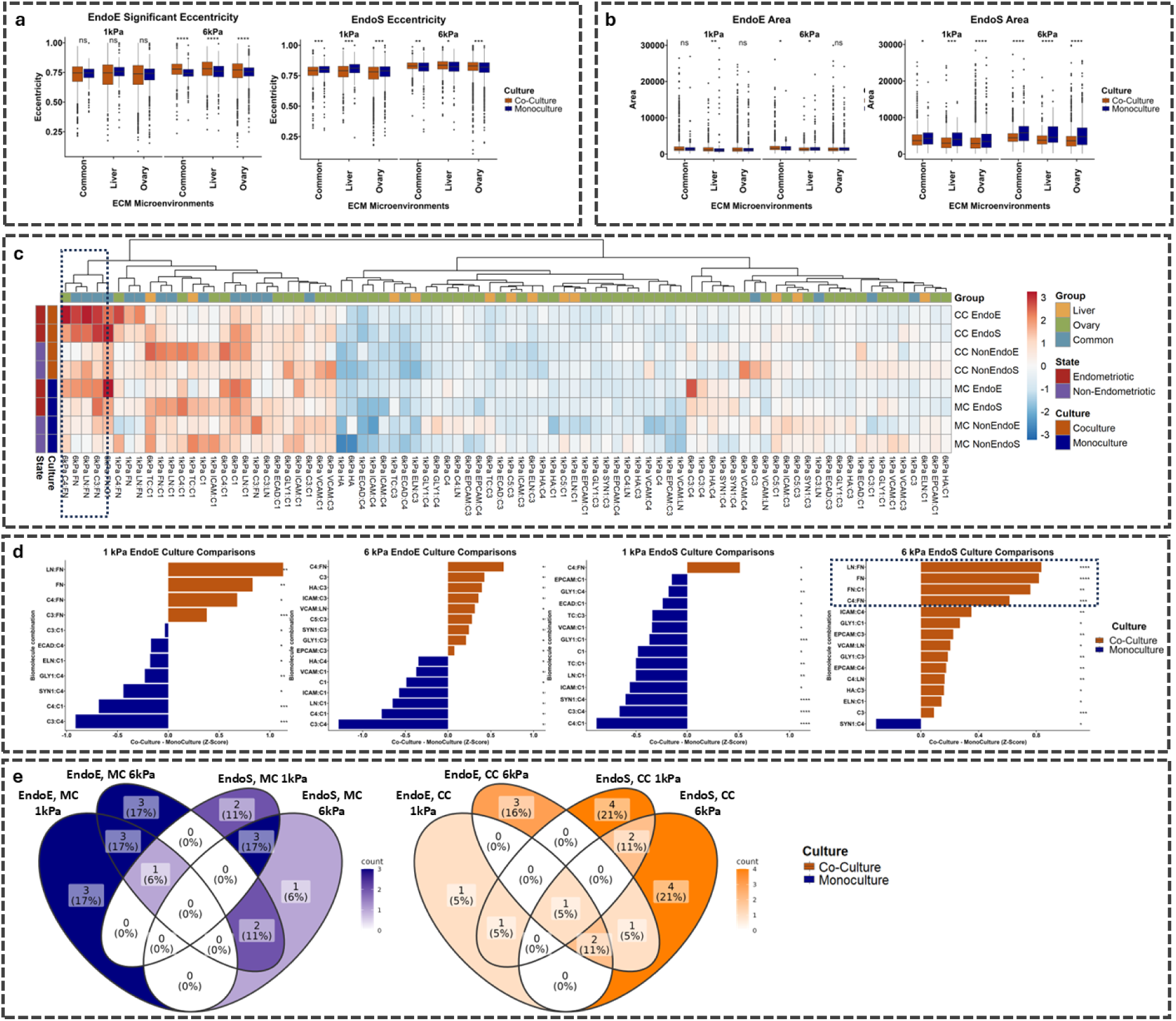
Coculture arrays examining cell eccentricity, area, and attachment differences. a) Quantification of eccentricity differences of EndoE and EndoS cells at both stiffnesses comparing coculture and monoculture grouped by tissue type. b) Comparison of coculture and monoculture area changes for EndoE and EndoS cells across stiffnesses and tissue type generalization. c) Heatmap comparing z-scores of attachment numbers for all four cell types across stiffnesses, culture conditions and grouped by tissue type. d) Significant differences in attachment z-score for EndoE and EndoS cells between coculture and monoculture conditions. e) Venn diagram of similarities in attachment conditions between coculture and monoculture for both endometriotic cell types.

When comparing cell attachment on soft (1 kPa) substrates (**Fig. 6c-d**), endometriotic epithelial cells in co-culture attached significantly more to common (LN:FN, FN, and C3:FN) and one ovary-specific (C4:FN) condition than in monoculture. On stiff (6kPa) islands, we observed significantly higher adhesion of endometriotic epithelial cells in co-culture vs. monoculture on all groups, but especially the ovary-specific group (C4:FN, HA:C3, ICAM:C3, VCAM:LN, SYN1:C3, GLY1:C3 and EPCAM:C3) (**Fig. 6d, S7e-h**). Endometriotic stromal cells responded differently than the endometriotic epithelial cells in co-culture. Co-cultured EndoS cells showed significantly higher attachment vs. monoculture on 1 kPa C4:FN (ovary) substrates (**Fig. 5d, S7c**). At 6 kPa, we saw a significant increase in endometriotic stromal attachments in co-culture over mono-culture on all groups, but generally on ovarian-specific (C4:FN, ICAM:C4, GLY1:C3, EPCAM:C3, HA:C3, C4:LN, VCAM:LN and EPCAM:C4) conditions (**Fig. 5d, S7d**).

Of note, there are ovary-specific conditions (C4:FN, HA:C3, VCAM:LN, EPCAM:C3 and GLY1:C3) where both endometriotic stromal and epithelial cells had significantly higher attachment on stiffer 6 kPa substrates. When we look at the adhesion profiles for all cell types in monoculture and co-culture, we observe that the adhesion profiles of endometriotic stromal and epithelial cells in co-culture are the most similar (**Fig. 6c, d**). Additionally, the top 4 conditions with significantly higher attachment of endometriotic stromal cells seem to have been increased to more closely match the adhesion profile of endometriotic epithelial cells (dotted box on **Fig. 6c, d**). This suggests that crosstalk between endometriotic stromal and epithelial cells improves endometriotic stromal cell attachment to matrix environments that already had high endometriotic epithelial cell attachment. To describe similarities in attachment profiles, Venn diagrams were generated to illustrate shifts in attachment commonalities between EndoE and EndoS at two stiffnesses in both monoculture or coculture (**Fig. 6e, Table S4**). These Venn diagrams include island conditions that were significantly higher in the endometriotic cells over their non-endometriotic counterparts (darker color indicates more ECM combinations, **Fig. 6e**). These results display shifts in attachment combinations from largely epithelial in monoculture to more stromal in coculture. Taken together, we observed significant differences in cellular attachment between disease cohorts in monoculture and coculture that emphasize the importance of cell signaling on attachment.

We also observed spatial patterns of epithelial and stromal cell attachment within the adhesive island as a function of disease state and ECM combination. We quantified this spatial patterning by describing the spatial distribution of stromal cells on each island as a function of radius (with dimensionless r = 0 indicating the center of the island and r = 1 indicating the edge of the island; **Fig. S8a**). Overall, we observed that non-endometriotic islands had higher stromal attachment than the endometriotic conditions over both stiffnesses (**Fig S8b**). Additionally, when separated into tissue groups, the stromal profiles of the endometriotic islands were not significantly different on the 1 kPa substrates, but on 6 kPa, the liver tissue group had a significantly higher stromal component than the other groups in the periphery of the island (r = 0.5 -1, **Fig. S8c**). Notably, LN:C1 and FN:C1 are two ECM combinations from the common group that showcase different stromal composition patterning (**Fig. S8d-g**). Here, while non-endometriotic stromal cells are more evenly distributed across the stiffer (6kPa) islands, we see increased attachment of endometriotic epithelial cells (reduced endometriotic stromal cell fraction) with endometriotic stromal cells more likely to partition to the periphery of islands rather than the center. This work represents the first profiling of multicellular cohorts on these islands and demonstrates the influence of stiffness, disease state, and ECM composition on patterning of cellular adhesion.

## 3. Discussion

Here, we report the development of a microarray-based system to investigate processes of endometrioma lesion initiation driven by selective attachment of endometriotic stromal and epithelial cells to the ovarian matrix. PA microarrays have been extensively used to examine the role of ECM on cell attachment; however, they had not been used to examine the role of cell adhesion molecules (CAMs, regulating cell-cell, or cell-matrix attachment) or cadherins (regulating cell-cell attachment). Studies have shown a complex interplay between cadherins and other ECM proteins regulating cell migration [46]. After identifying several CAMs and cadherins of interest (VCAM-1 [54], ICAM-1 [55], EP-CAM [56], E-cadherin [57]), we confirmed their bioactivity in a 2D array format – the first-time complex adhesion motifs involving both CAMs and conventional matrix proteins and GAGs were deployed together in this array platform. We observed minimal cellular attachment to islands containing solely CAMs or cadherins, regardless of substrate stiffness (1kPa and 6kPa) or the inclusion of calcium for cadherin activation (Ca; 10 nM, 5 nM, 2.5 nM, 0 nM). Interestingly, we observed improved adhesion for CAMs or cadherins when printed with an additional ECM molecule (e.g., fibronectin), which supported our decision to include these biomolecules for further analysis. To facilitate our analyses, after performing a first-round comprehensive screen, we removed lower adhesion combinations (<25 cells / island average) to bolster our work by increasing the number of islands per biological replicate. This reduction in combinations resulted in 42 combinations of interest (**Table 2**).

Analysis of cell attachment patterns revealed increased eccentricity in endometriotic cells in comparison to their non-endometriotic derived counterparts (**Fig. 3c**), aligning with what researchers consider to be disease-state cells, especially those undergoing EMT [47]. Cell shape may play a significant role in signal transduction, including downstream kinase activity [48]. Receptor tyrosine kinases (RTKs) have been implicated in endometriosis persistence as they command differentiation and metabolism of cells and are found in increased rates in endometriotic lesions and are being investigated as potential therapeutic targets for endometriosis treatment [49]. Atypical mechanosensing in relation to lack of cell rigidity-sensing complexes can result in untamed growth (like in cancer) [27]. Investigators found that decreased substrate stiffness, more attuned to physiologically relevant stiffness of the ovaries, decreased f-actin stress fiber formation *in vitro*, and that increased substrate stiffness favored stromal cell proliferation [20]. Although both endometriotic cells displayed higher eccentricity than their non-endometriotic counterpart, only endometriotic stromal cells were significantly larger than non-endometriotic stromal cells. Endometriotic epithelial cells in monoculture had a significantly smaller area compared to all other cell types (**Fig. 3e**). We also observed a much wider distribution of cell area for the stromal populations (**Fig. 3e & S2b**). Cell paracrine and juxtocrine signaling could explain this discontinuity between the relationship between endometriotic epithelial and stromal cell relationship with stiffness, which these previous studies have not yet considered.

Endometriotic epithelial cell attachment patterns were modulated most significantly more in response to substrate stiffness and ECM composition compared to non-endometriotic epithelial cells (**Fig. 4b-d**). Endometriotic epithelial cells attached in higher numbers to the stiffer substrates. When comparing our two stiffness conditions to disease state, the 6 kPa environment that the endometriotic epithelial cells favored is considered more similar to an older or more fibrotic ovary. Collagen IV, a component of all three ECM combinations (C3:C4, C4:C1, and C4:FN), that regardless of substrate stiffness, endometriotic epithelial cells significantly attached to over their non-endometriotic complement, is an integral component of the ovarian ECM as well as a soluble factor released during ovulation [4, 50]. The endometriotic stromal cells were also altered more in relation to ECM composition than non-endometriotic derived stromal cells (**Fig. 5c-d**). Here, however, we do not see significant differences in attachment number based on stiffness (**Fig. 5b**). Previous ECM attachment studies on endometriotic cell populations have shown that endometriotic cells are generally more adhesive than non-endometriotic derived cells, which aligns with our findings [4, 51].

The most significant ECM signals on stiffer (6 kPa) substrate that increased endometriotic epithelial cell attachment were Collagen I and Fibronectin. Besides being the most common collagen in the body, collagen I has been implicated in the advancement of endometriosis through endometriotic myofibroblast production leading to fibrosis [20]. A genome-wide study associated the gene FN1, which regulates the production of fibronectin, with severe states of endometriosis [20] alongside variances in FN between non-endometriotic and endometriotic patients [52–54]. Collagen I and Fibronectin also promoted higher attachment for endometriotic vs. non-endometriotic stromal cells across various matrix combinations, including C4:C1 (1 kPa), LN:C1 (6 kPa), FN:C1 (6 kPa), and C3:FN (6 kPa) (**Fig. 4H**), as well as laminin. Laminin-1 also played a significant role in endometriotic stromal cell attachment, consistent with findings that laminin is found in high levels in the serum of women with late-stage endometriosis [55, 56]. Laminins function as a major component of basement membranes (BMs), and they play an integral role in cell adhesion and migration [57]. It is intriguing that endometriotic cells are attaching at higher rates to some combination including Collagen III than that of non-endometriotic derived cells, given that Collagen III is expressed in higher levels in the native endometrium than Collagen I [58]. However, increased deposition of collagen III has been associated with various types of solid tumor fibrosis [59, 60]. Ongoing work is exploring the study of these motifs in 3D hydrogel models in the presence or absence of endothelial cells. Examining these combinations in combination with epithelial cells in order to provide additional insight regarding mechanisms governing increased attachment.

When examining the role of ECM on spatial patterning of endometriotic cells, we observed conditions lending to stromal cell attachment especially along the edges of the islands. Island edges that saw stromal cell preference could be compared to extensions of the ovary called the corpus luteum (CL). The CL is considerably stiffer than the surrounding matrix, and requires intense remodeling and degradation after ovulation [61]. Researchers have also recently found that patients could experience endometriomas at the site of ovulation, so the preferential cell attachment at the boundary edges, on stiffer substrates, could indicate this is a preferential site for early attachment [62].

These studies aimed to investigate the mechanism by which early endometriotic lesions attach, specifically investigating the potential (tropism) preferences of cellular attachment. Five conditions that endometriotic epithelial cells attached more significantly to than their healthy complement (NonEndoE) on 1 kPa substrates were ovarian specific ECM combinations (the remaining two were common ECM combinations). Higher stiffness (6 kPa) condition results were less conclusive, with 4/10 significantly higher diseased attachment conditions being attributed to the ovaries (and the remainder being attributed to common ECM combinations) (**Fig. 4d**). We observe a similar trend with endometriotic stromal cells; 5/6 of the ECM combinations that EndoS in monoculture at 1 kPa attached more significantly to NonEndoS were ovarian labeled; at 6 kPa, this ratio is 3/6 ECM conditions (**Fig. 5d**). The argument for ovarian tropism is less compelling in epithelial-stromal cocultures when considering endometriotic vs. non-endometriotic epithelial attachment (1 kPa = 1/5 conditions of increased attachment were ovary; 6 kPa = 3/7 ovary) (**Table S5**). However, we regain the significance of ovarian tropism when considering endometriotic stromal cells in co-culture (EndoS: 1 kPa = 6/8 ovary; 6 kPa = 5/10 ovary). If we look globally at ECM conditions where endometriotic epithelial cell attachment is more significant in co-culture vs. monoculture, we can revisit tropism with 7/9 of the 6kPa ECM combinations being ovarian-specific. For endometriotic stromal cells, the only significant attachment at 1kPa in coculture was ovarian-specific and at 6kPa 9/14 were ovarian. Collectively, these data suggest endometriotic cells are more significantly influenced by tissue type and composition on lower-stiffness substrates. On stiffer substrates that are more consistent with fibrotic lesions, we did not observe definitive tropism to ovarian attachment factors in monoculture. These findings may have significance in the context of understanding initial lesion initiation versus lesion recurrence after ablation, where post-ablation lesions would be more likely to recur due to the increased stiffness due to fibrosis on the microenvironment [63]. In co-culture, however, we observe higher tropism to ovarian ECM on both endometriotic cell types [64]. Recently, researchers have investigated the role of stiffness on endometriotic epithelial cell morphology and phenotype, finding these cells maintained a rounded morphology at 2 kPa but elongated at 30 kPa, however they also observed that a stiffer matrix potentially created a positive feedback loop causing further fibrotic changes in endometriotic cells via increased proliferation, collagen I deposition, and differentiation [20, 63, 65]. Here, one potential mechanism driving this positive feedback loop of fibrosis is the mechanosensitive YAP/TAZ pathway [66], inspiring ongoing experimental efforts using these array systems.

Changes in cell area and morphology in endometriotic stomal cells in coculture suggest they may be taking on a different fibroblast phenotype. Cell traction forces are positively correlated with increased cell area [67], with myofibroblasts, often associated with fibrosis in endometriosis, typically larger than normal fibroblasts [68]. Previous studies have found that endometriotic epithelial cells undergo a partial EMT-process when cultured for extended periods on stiffer substrates resulting in an even further increase in fibrotic environments [20, 69]. Prior studies examining the role of substrate stiffnesses on endometrial adenocarcinoma-derived epithelial (Ishikawa) cells found them to be resistant to changes in stiffness up to 23 kPa [70]. This suggests an innate difference in mechanosensing between the endometriotic cells and their non-endometriotic complements. Perhaps non endometriotic epithelial cells are more resistant to changes in stiffness across the range we tested (1 kPa vs. 6 kPa), as the native endometrium undergoes rapid cycling during the menstrual cycle. Taken together, our findings suggest cohorts of endometriotic cells function as a unit, attaching more similarly in coculture than in monoculture. We also corroborate previous findings suggesting endometriotic cells seeded on stiffer substrates function in a pseudo positive-feedback loop developing a more fibrotic morphology (more elongated) and size (myofibroblasts are larger compared to fibroblasts). These cells may exacerbate the fibrotic environment by depositing more matrix and through secreted proteins [71]. We also observed increased modulation of cell attachment at lower stiffness substrates, which supports the notion of the importance of early ovarian tissue tropism on the activity of endometriotic cells. Although this study was conducted in 2D, we hypothesize that the effects of cell attachment molecules intrinsic to the ovary that enhanced endometriotic attachment will also be imperative to invasion in 3D models. Recently, our group has described the use of high-throughput (2D) matrix array systems to as evaluate cell chirality and translate findings to a well-characterized gelatin hydrogel (3D) model of the endometrial tissue microenvironment [38]. These platforms provide a framework to investigate the influence of heterogeneous cell-matrix and cell-cell interactions in the context of endometriotic cell activity. Most importantly, our findings clearly support the hypothesis that to properly recapitulate the complex tissue architecture during endometrioma lesion initiation, we must develop better 3D models that incorporate multicellular components, account for the stiffness of the tissue (especially based on disease status and age), and account for the variation in extracellular matrix and attachment ligands.

## 4. Materials and Methods

### 4.1. Cell Culture & Maintenance

To represent the native, eutopic endometrium (NonEndoE & NonEndoS respectively), primary endometrial epithelial cells, EECs, (LifeLine Cell Technology, FC-0078), and T-human endometrial stromal cells, ESCs, (ATCC, CRL-4003) were cultured. We chose to represent endometriotic lesions (EndoE & EndoS respectively) with immortalized human endometriotic epithelial-like cells, 12z, (ABM, T0764) and immortalized ectopic endometrial stromal cells, iEC-ESCs, kindly provided by Dr. Asgi Fazleabas. EECs, ESCs, and 12z cells were cultured according to manufacturer’s specifications, substituting charcoal-stripped fetal bovine serum (Sigma-Aldrich, F6765) in place of traditional FBS. iEC-ESCs were cultured in phenol-red free DMEM/F-12 with 1mM sodium pyruvate (SCS Cell Media Facility, UIUC), supplemented with 10% charcoal-stripped fetal bovine serum (Sigma-Aldrich, F6765), and 1% penicillin/streptomycin (Gibco, 15-140-122). Cells were cultured in a 37°C incubator with 5% CO_2_. All cells were routinely tested for mycoplasma to confirm cell quality and lack of contamination (Tumor Engineering and Phenotyping Shared Resource, UIUC).

### 4.2. Polyacrylamide Microarray Fabrication

Polyacrylamide (PA) hydrogels were prepared as previously described [37–39, 41]. Briefly, 25 mm x 75 mm glass microscope slides were washed, etched with 0.2 N NaOH, and silanized with 2% v/v 3-(trimethosy silyl) propyl methacrylate in ethanol. Prepolymer 1kPa and 6kPa PA solutions with defined elastic moduli determined by acrylamide/bisacrylamide w/v ratios and similar porosity were fabricated [39, 72]. 1 kPa solutions were comprised of 4% acrylamide (Sigma-Aldrich, A3553-100G) and 0.4% bis-acrylamide (Sigma-Aldrich, M7279-25G) and 6_JkPa solutions were made of 6% acrylamide and 0.45% bis-acrylamide as described previously [39, 72]. The prepolymer solutions were then mixed with 20% w/v solution of Irgacure 2959 (BASF, 50047962) in methanol to achieve a final solution ratio of 9:1 (prepolymer to Irgacure). This final solution was deposited onto the silanized slides and contained with 22 mm × 60 mm coverslips, then exposed to 365 nm UV A light (240 × 10^3^ μJ) for 10 minutes. After UV polymerization, the cover glass was removed, and the slides were washed in dH_2_O for 3 days at room temperature.

Microarray fabrication was completed as previously described [39, 72]. The hydrogels were dehydrated on a hot plate for ≥15 minutes at 50_J°C. ECM and attachment factor biomolecules (Table 1 & 2) [73–81] were diluted in 2x ECM printing buffer (40% v/v glycerol, 0.5% v/v warmed Triton-X, and 0.8% v/v glacial acetic acid in dH_2_O with 16.40 mg/mL sodium acetate and 3.72 mg/mL EDTA) to a final working concentration of 250 μg/mL and loaded into a 384-well V-bottom microplate (USA Scientific, 1823-8400) using a robotic liquid handler (OT-2, Opentrons). The microarrayer (OmniGrid Micro, Digilab) was loaded with SMPC Stealth microarray pins (ArrayIt, SSP015) to microprint ECM combinations into ∼600 μm diameter islands. It was experimentally determined that 2 mM concentration of CaCl_2_ added to the initial protein solution was adequate for cell attachment without over-diluting the cadherin of interest (**Fig. S1a**).

### 4.3. Microarray Experimentation

Microarrays were sterilized with 1% penicillin/streptomycin (Gibco, 15-140-122) in phosphate buffered saline (PBS, Sigma-Aldrich D5773-10L) and UV light for 30 minutes in a biological safety cabinet. For coculture experiments, stromal cells were treated with CellTracker™ Green CMFDA Dye (ThermoFisher, C7025) in a 3:1000 ratio (CellTracker™:Media) for 20 minutes at 37°C. Similarly, epithelial populations were treated with CellTracker™ Orange CMRA Dye (ThermoFisher, C34551) in a 1:200 ratio (CellTracker™:Media) for 20 minutes at 37°C. Cells were centrifuged at 250 rcf for 5 minutes, washed with PBS, and centrifuged again at 250 rcf for 5 minutes. Arrays composed of co-cultured cells in a 1:3 ratio of epithelial to stromal based on literature [34, 82]. Cells were passaged and seeded at a density of 200,000 cells per microarray (for monoculture experiments) in 500 μL of respective complete culture medium, or 50:50 media for coculture experiments.

After a 30-minute incubation at 5% CO_2_ and 37°C, each slide was washed with sterile PBS, followed by gentle addition of 4 mL of respective complete culture before incubating at 5% CO_2_ and 37°C for 24 hours. For each set of experiments, 2-3 biological replicates with temporally separate printing and seeding experiments were performed. Within each biological replicate, each combination of stiffness and cell type were replicated on three slides, each with 5-10 islands per condition.

### 4.4. Immunostaining & Mounting Slides

After 24 hours of incubation, the microarrays were washed (3x) with PBS. Then, arrays were fixed (1x) with formalin for 15 minutes, quick washed (3x) with PBS, permeabilized (1x) with 0.5% Tween20 for 15 minutes, quick washed (3x) with PBST (50 μL Tween 20, 50mL PBS) for 5 minutes each, and then blocked (1x) with 2% Abdil (1g BSA, 50 μL Tween 20, 50mL PBS) for 1 hour. The microarrays were stained with phalloidin (abcam, ab176758) diluted in 2% abdil (1:1000 respectively) covered from light for 1 hour. The microarrays were quickly washed (3x) with PBS. After the washes, the PBS was removed from the slides, 100 μL of DAPI Fluoromount-G (SouthernBiotech, 0100-20) solution was added on top of the microarrays and sandwiched with a coverslip (Electron Microscopy Sciences, 63765-01), sealed with clear nail polish, and stored protected from light at 4°C until imaging. All steps were performed at room temperature unless otherwise specified.

### 4.5. Microarray Imaging & Analysis

Cellular microarrays were imaged at 10x and 20x magnification on the Zeiss AxioScan.Z1 Slide Scanner. Images of entire arrays were converted to 8-bit TIFF files per output channel using Fiji (ImageJ) and cropped into single island images utilizing dextran-rhodamine marker islands to identify the location of arrayed conditions within each image. CellProfiler (version 4.2.6) was used to identify and measure nuclei and fluorescent regions using the IdentifyPrimaryObjects, IdentifySecondaryObjects, and MeasureObjectSizeAndShape modules [83]. Single-cell fluorescent intensity was quantified using the MeasureObjectIntensity module. The output from CellProfiler was loaded into RStudio for analysis. Cell location for patterning analysis was determined using coordinates output by CellProfiler. Cell location was assigned based on distance from the centroid [39, 84]. For co-culture analysis, the fluorescent intensity of the CellTracker dyes were rescaled to stretch each image to full intensity range using RescaleIntensity module and then binned for the expression of the pink and green channels by mean intensity using ClassifyObjects. Additionally, the comparison of mean intensity between the pink and green channels were classified as low-low, low-high, high-low, and high-high for mean intensity using the ClassifyObjects module. These classifications were used to create a decision tree that reliably assigned 95.41% of the total cells as either Stromal or Epithelial, with the remaining cells being categorized as Unassigned due to not fitting the criteria of the decision tree. For analysis, the number of cells attached to each island were counted and individual cell measurements were averaged together. Each representative island for a condition on a slide were averaged together to provide a per slide measurement. Each island serves as a technical replicate of a condition on a slide. The multiple slides per experimental run serve as technical replicates of the culture conditions. For Venn diagram analysis of groups, we utilized ggVennDiagram (ShinyApp) [85].

### 4.6. Statistics

All array experiments consisted of at least three biological replicates, unless otherwise specified. For each biological replicate, 30 technical replicates (total individual islands per ECM condition) were presented over 3 slides, unless otherwise specified. For comparisons between cell types, z-scores were calculated by subtracting the cell type mean from the averaged response and dividing that cell type standard deviation for each biological replicate. Linear regression analyses were performed using the lmer() function from the lme4 package in R [86]. The outputs of each regression are presented as coefficient estimates and associated standard error. The marginal and conditional R^2^ value as well as the intraclass correlation are provided for each model, as calculated by the tab_model function from sjPlot R package [55]. The MASS package [56] was used to perform Box Cox analysis to identify appropriate response transformations for better model fit. For comparisons between conditions, non-parametric Wilcoxon tests were performed using the compare_means and stat_compare_means functions from the ggpubr package in R [57]. P values of <0.05 were considered significant for all statistical comparisons. (* p ≤ 0.05, ** p ≤ 0.01, *** p ≤ 0.001, **** p ≤ 0.0001).

## Acknowledgements

The authors would like to thank Austin Cyphersmith (IGB Core Facilities) for technical guidance in high-content imaging and analysis. Funding sources include the National Institute of Diabetes and Digestive and Kidney Diseases of the National Institutes of Health under Award Number R01 DK099528 (BACH) and R01 DK125471 (GHU); the National Cancer Institutes of the National Institutes of Health under Award Number R01 CA256481 (BACH); National Institute of Biomedical Imaging and Bioengineering of the National Institutes of Health under Award Number T32EB019944 (HRCK); and the Eunice Kennedy Shriver National Institute of Child Health and Human Development of the National Institutes of Health (T32 HD108075 (HST). The content is solely the responsibility of the authors and does not necessarily represent the official views of the NIH. The Center for Gender & Sex in Health is supported a Strategic Research Initiative award from the Grainger College of Engineering at the University of Illinois. The authors are also grateful for additional funding provided by the Department of Chemical & Biomolecular Engineering, the Carl R. Woese Institute for Genomic Biology, the Cancer Center at Illinois, and the Illinois Scholars Undergraduate Research (ISUR) Program at the University of Illinois Urbana-Champaign for funding Sarah Hashim as an undergraduate researcher.

## Contributions (CRediT: Contributor Roles Taxonomy [87, 88])

**H.S. Theriault:** Conceptualization, Data curation, Formal Analysis, Visualization, Investigation, Methodology, Writing – original draft, Writing – review & editing. **H.R.C. Kimmel:** Conceptualization, Data curation, Formal Analysis, Visualization, Investigation, Methodology, Writing – original draft, Writing – review & editing. **A. Nunes:** Investigation, Methodology. Writing – review. **A. Paxhia:** Investigation, Methodology, Writing – review. **S. Hashim:** Investigation. **K.B.H. Clancy:** Conceptualization, Methodology, Writing – review & editing. **G.H. Underhill:** Conceptualization, Methodology, Writing – review & editing, Funding acquisition. **B.A.C. Harley:** Conceptualization, Resources, Project administration, Funding acquisition, Supervision, Writing – review & editing.

