## Supplemental Figures 1-6 for "Matrix tropism influences endometriotic cell attachment patterns"

**Hannah S. Theriault**^1,2*^**, Hannah R.C. Kimmel**^1,2*^**, Alison C. Nunes**^2^**, Allison Paxhia**^1^**, Sarah Hashim**^3^**, Kathryn B.H. Clancy^2,4,5,6^, Gregory H. Underhill**^1,2^**, Brendan A.C. Harley**^2,3,6,7^

^1^ Dept. of Bioengineering

^2^ Carl R. Woese Institute for Genomic Biology

^3^ Dept. Chemical and Biomolecular Engineering

^4^ Dept. of Anthropology

^5^ Dept. of Women & Gender Studies

^6^ Center for Gender and Sex in Health

^7^ Cancer Center at Illinois

University of Illinois at Urbana-Champaign

Urbana, IL 61801

**Corresponding Author:**

B.A.C. Harley

Dept. of Chemical and Biomolecular Engineering

Cancer Center at Illinois

Carl R. Woese Institute for Genomic Biology

University of Illinois at Urbana-Champaign

110 Roger Adams Laboratory

600 S. Mathews Ave.

Urbana, IL 61801

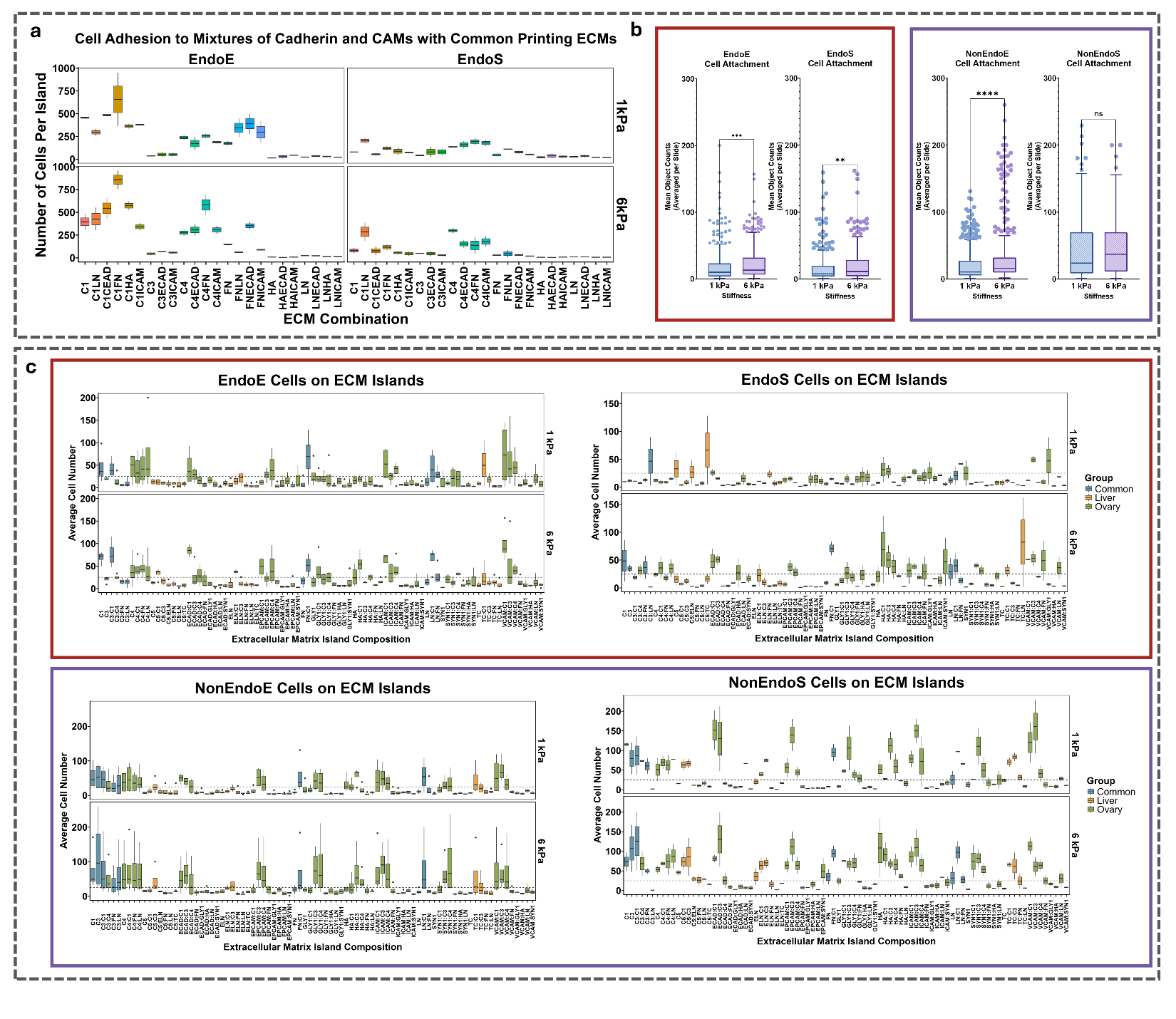
 **Figure S1: Extended figures for examining the ability of PA Microarrays to support the addition of CAMs and Cadherins.** **a)** EndoE and EndoS cells were seeded with some preliminary mixtures of common ECM components, E-cadherin and ICAM. Results are displayed as the number of cells per island. **b)** On the full microarray with all combinations, we observed significant differences in attachment at 6 kPa > 1kPa for all cell types except NonEndoS. **c)** Boxplots comparing the average cell number of each cell type over 1 kPa and 6 kPa and each ECM combination described by tissue type grouping.

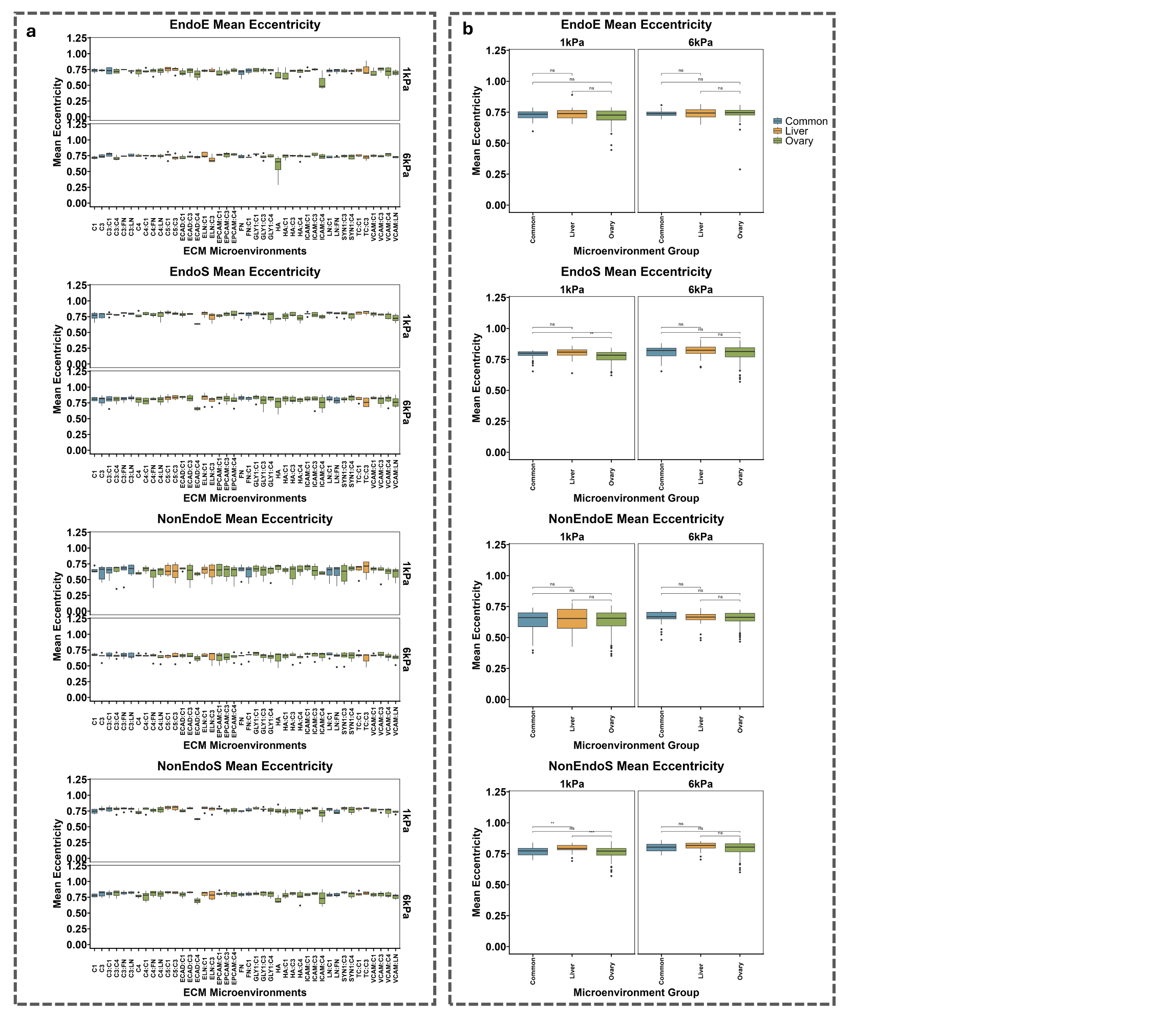
**Figure S2: Reduced arrays eccentricity results for all cells.** **a)** Quantification of eccentricity of all four cell types split by stiffness and across all ECM combinations labeled by tissue type. **b)** Eccentricity for each cell type quantified averaged across each tissue category.

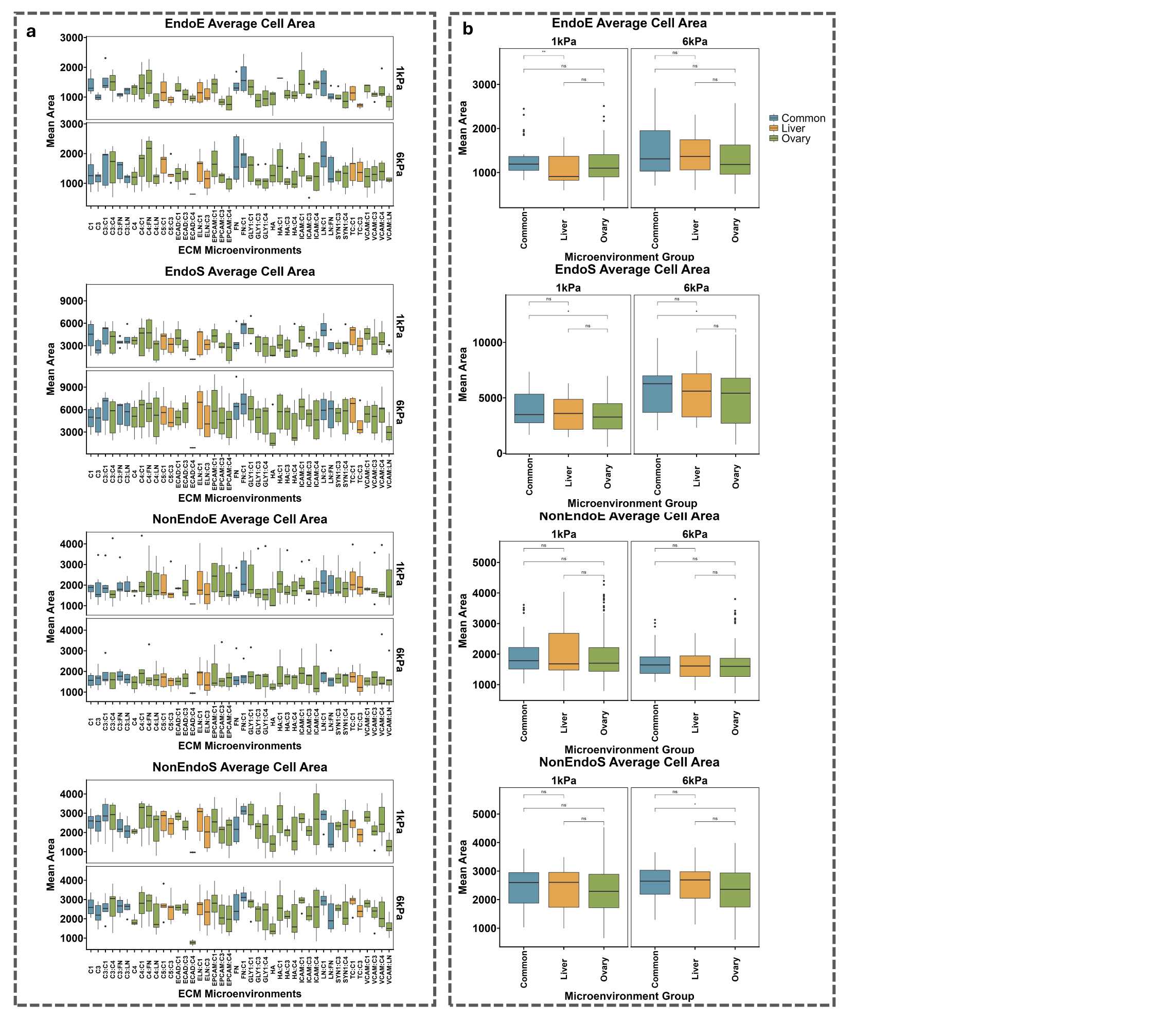
 **Figure S3: Reduced arrays area results for all cells.** **a)** Quantification of area of all four cell types split by stiffness and across all ECM combinations labeled by tissue type. **b)** Area for each cell type quantified averaged across each tissue category.

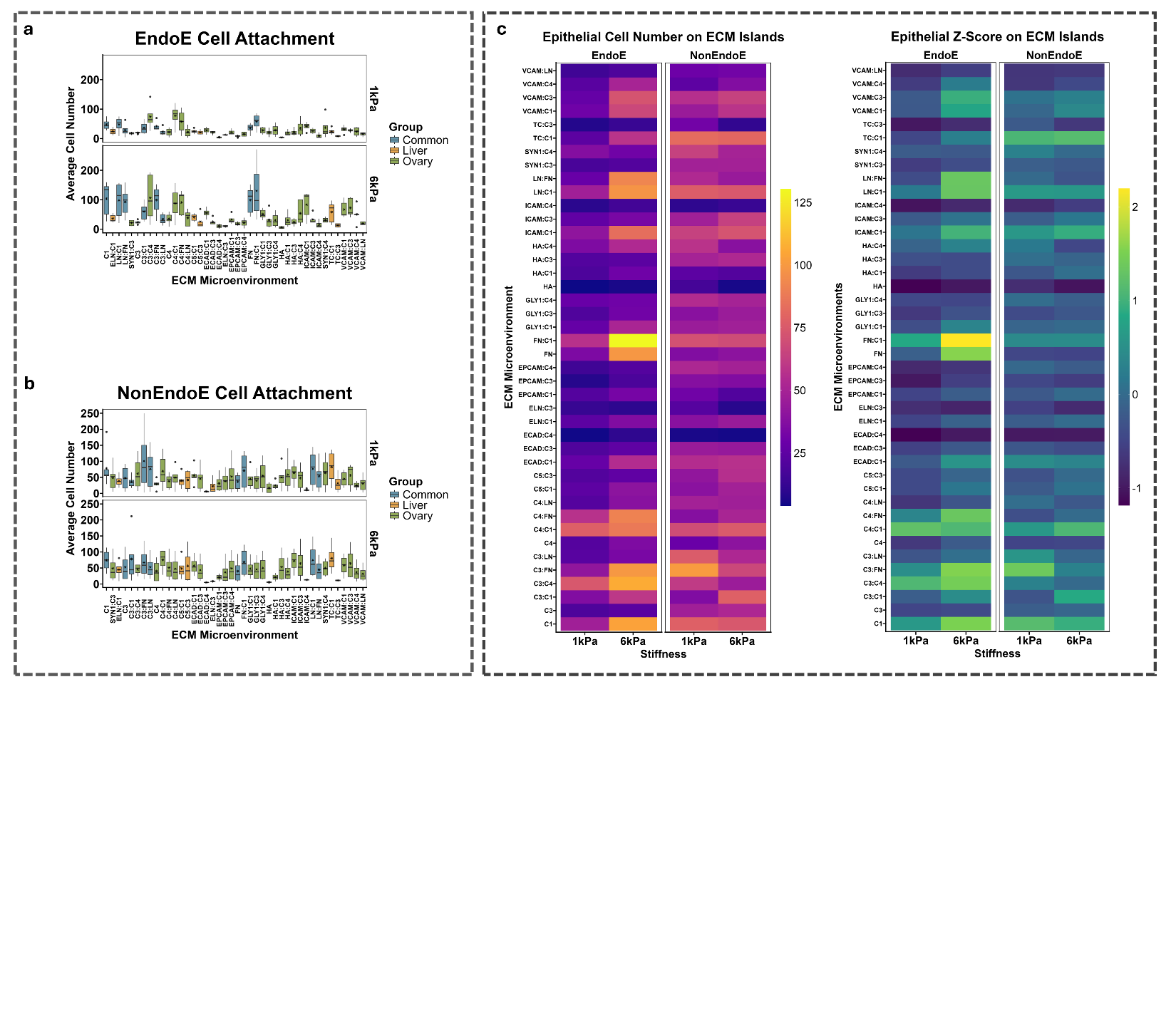
**Figure S4: Reduced arrays comparison of epithelial cell attachment extended data. a)** Quantification of the average cell number on each island condition for EndoE cells and **b)** NonEndoE cells. **c)** Heatmaps describing cell attachment comparing EndoE and NonEndoE cells alongside the z-score examination.

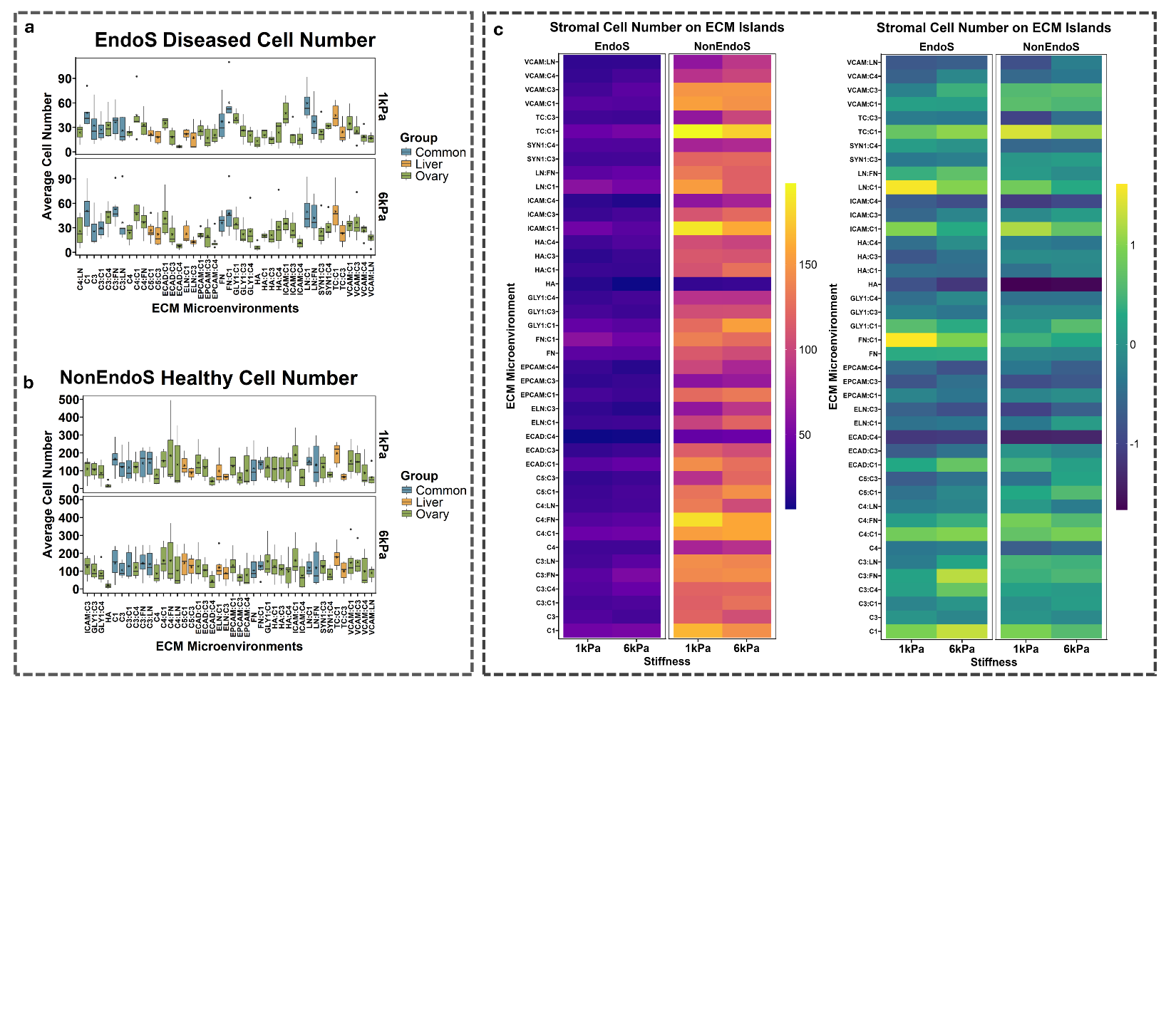

**Figure S5: Reduced arrays comparison of stromal cell attachment extended data. a)** Quantification of the average cell number on each island condition for EndoS cells and **b)** NonEndoS cells. **c)** Heatmaps describing cell attachment comparing EndoS and NonEndoS cells alongside the z-score examination.

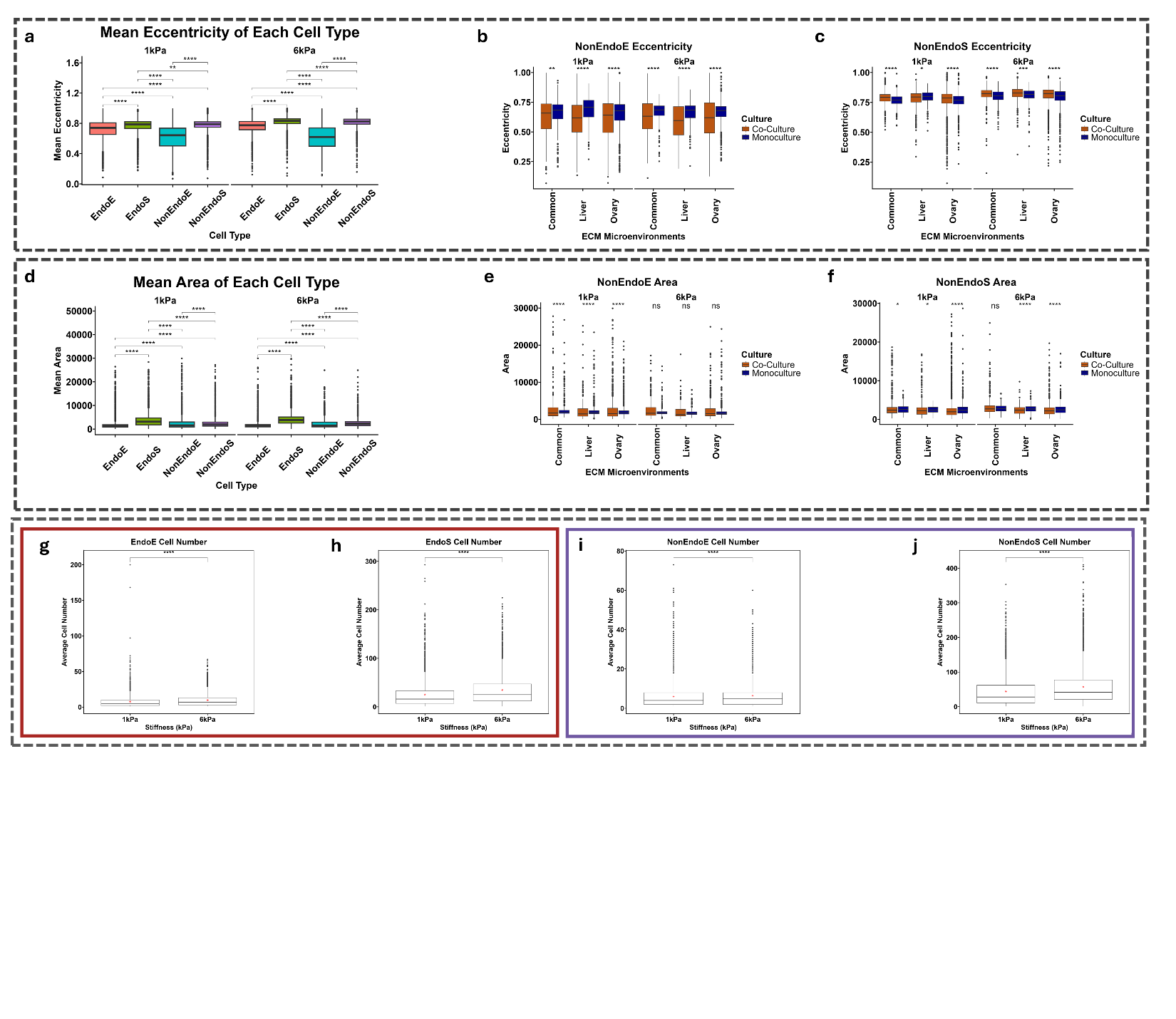

**Figure S6: Coculture arrays examining cell eccentricity and area differences.** **a)** Quantification of the mean eccentricity of each cell type across both stiffnesses in coculture. **b)** Comparison of coculture and monoculture eccentricity changes for NonEndoE and NonEndoS cells across stiffnesses and tissue type generalization. **c)** Quantification of the mean area of each cell type across both stiffnesses in coculture. **d)** Comparison of coculture and monoculture area changes for NonEndoE and NonEndoS cells across stiffnesses and tissue type generalization. **e)** Average cell number of each cocultured cell type based on stiffness.

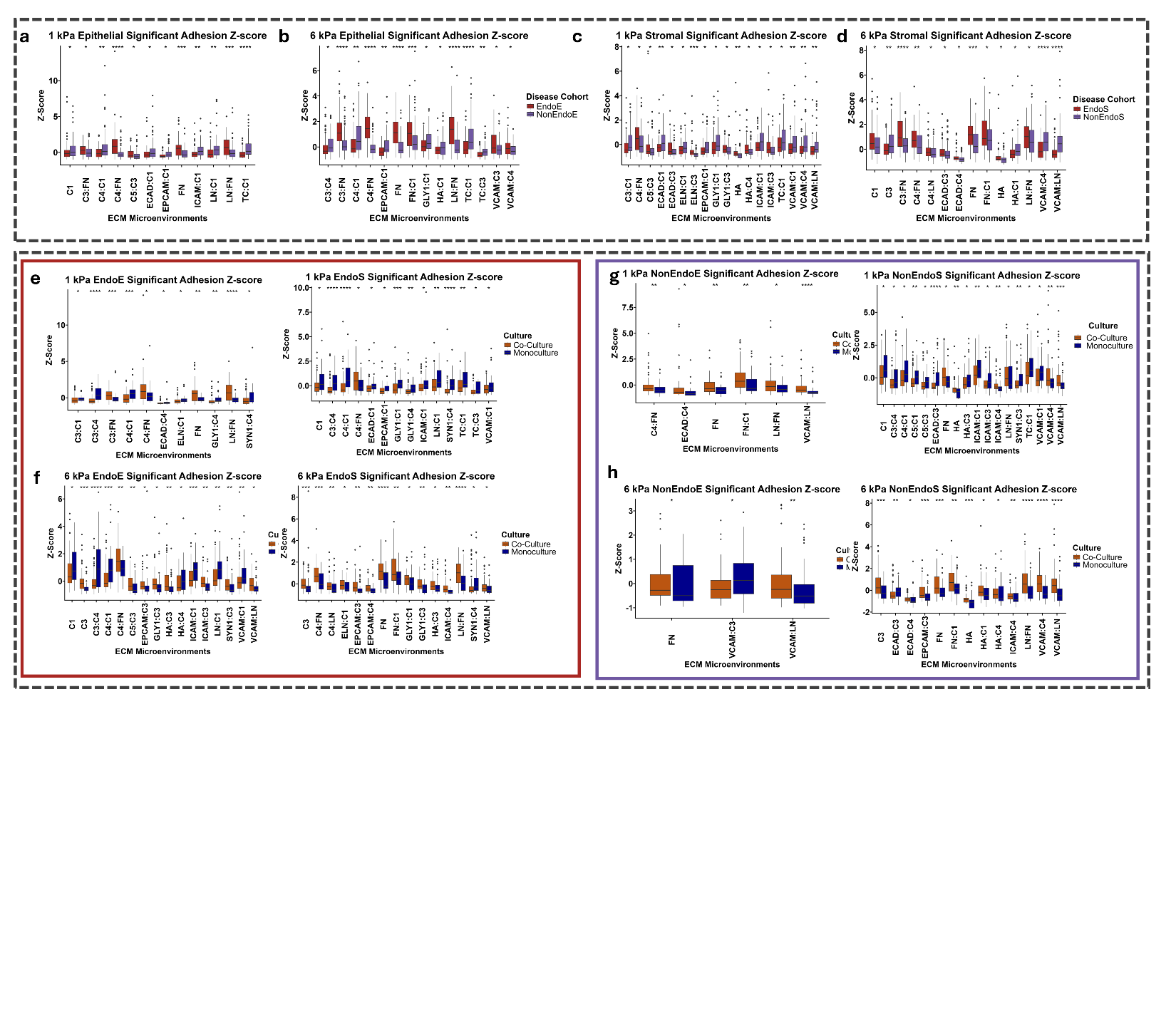

**Figure S6: Coculture arrays examining cell attachment differences.** **a)** Quantification of significantly different attachment z-scores of cocultured epithelial cells at 1 kPa and **b)** 6 kPa. **c)** Further quantification of significant differences in attachment z-scores for cocultured stromal cells at 1 kPa and **d)** 6 kPa. **e)** Significant z-score comparisons between EndoE and EndoS in coculture vs. monoculture at 1 kPa and **f)** 6 kPa. **g)** Significant z-score comparisons between NonEndoE and NonEndoS in coculture vs. monoculture at 1 kPa and **h)** 6 kPa.

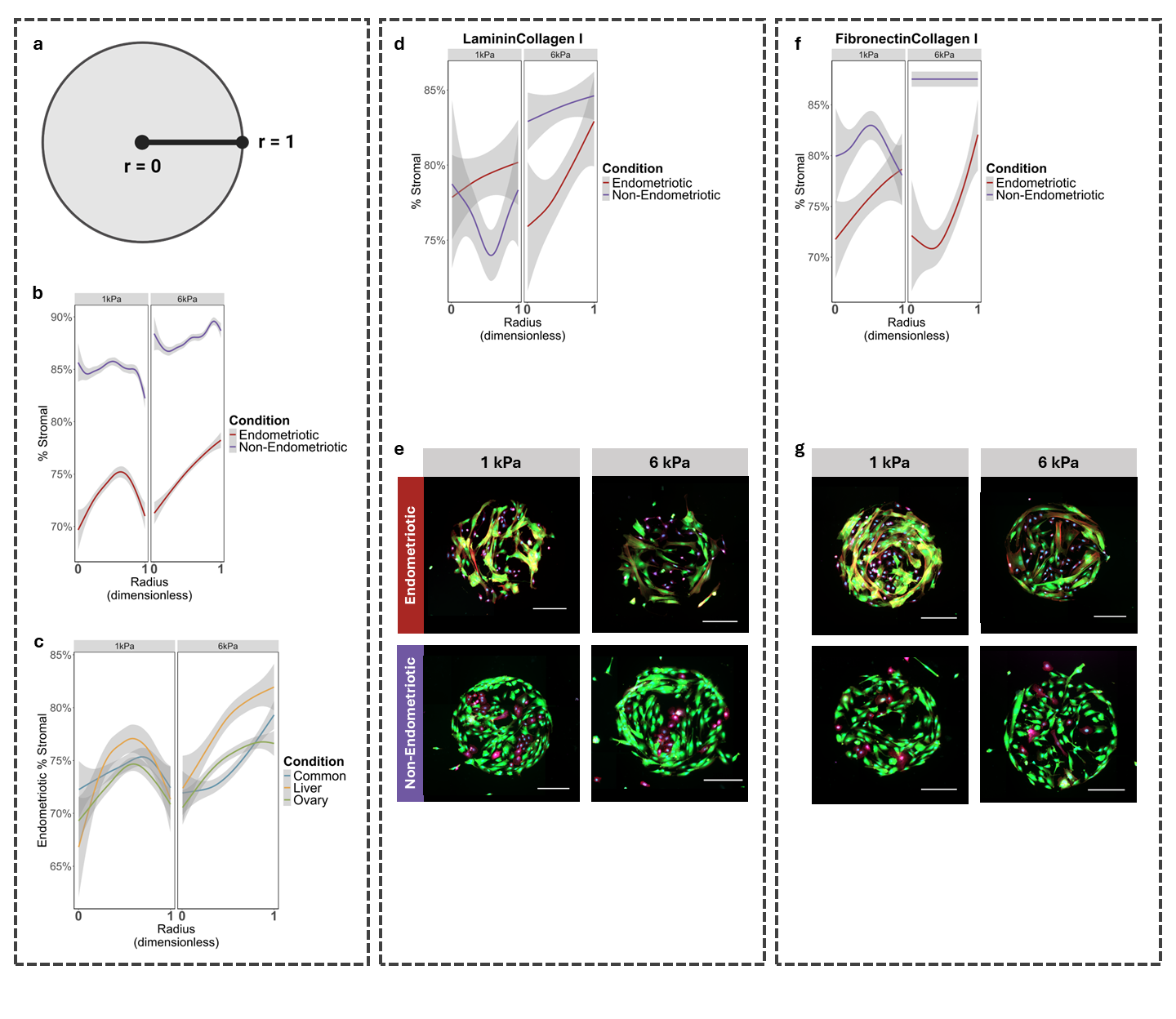

**Figure S6: Example spatial patterning of coculture arrays.** **a)** Spatial patterning on islands was quantified as the spatial distribution of stromal cells on each island as a function of the island radius. Indication for dimensionless measures of radius for each island with dimensionless r = 0 indicating the center of the island and r = 1 indicating the edge of the island. **b)**. Endometriotic islands over both stiffness conditions had significantly higher stromal cell attachment than the endometriotic islands. **c)** Endometriotic islands did not present significantly differential spatial patterning dependent on tissue group at 1kPa; however, at 6kPa, we observed significantly larger stromal contribution at approximately r = 0.5 for the liver condition over the ovarian and common tissue groups. **d)** Endometriotic and non-endometriotic cells on Laminin and Collagen I islands and **f)** Fibronectin and Collagen I islands. Representative islands of each condition are displayed in **e)** and **g).** Data were plotted as lines with 95% CI plotted in gray for spatial patterning. Scale bar indicates 200 μm.
